# Unveiling the control of N and P on DOM fate in a Mediterranean coastal environment

**DOI:** 10.1101/2024.03.20.585141

**Authors:** Clara Dignan, Véronique Lenoble, Chiara Santinelli, Giancarlo Bachi, Duc Huy Dang, Nicole Garcia, Benjamin Misson

## Abstract

Dissolved organic matter (DOM) and heterotrophic prokaryotes (HP) are key players in the oceanic carbon cycle. Although several biotic and abiotic factors controlling DOM fates are known, the hierarchy of their respective influences is still debated. Two contrasting Mediterranean coastal sites were sampled: a harbour under strong continental and anthropogenic influence (T) and an open coastal area (G). Interestingly, similar dissolved organic carbon (DOC) concentrations were observed in both samples. However, they showed marked differences in dissolved inorganic nitrogen and organic phosphorus concentrations (60-fold and 80% higher value in T), as well as in DOM optical properties and molecular composition. Incubation experiments were performed to expose the HP communities of each site to dissolved substances from T and G for three weeks. DOC removal was similar (−10 %) regardless HP origin and dissolved substances characteristics. HP growth and their maximal abundance were higher (+ 300 %) with dissolved substances from T, regardless HP origins. This indicates different fates of DOC processed by microbial communities as a function of abiotic determinants. Higher HP growth was associated to elevated initial content and higher consumption of inorganic nitrogen, organic phosphorus, three fluorescent DOM components, nitrogen-containing molecules and carbohydrates. These results provide insights into the main drivers of marine DOM fate: at similar DOC concentrations and low inorganic P concentrations. We evience the preferential consumption of lignin-like compounds where theoretically more labile molecules were available, thus reinforcing the need of in depth molecular studies for a better understanding of DOM-microbes interactions in the ocean.

## 1 Introduction

Dissolved organic matter (DOM) and heterotrophic prokaryotes (HP) are well recognized to be key players in the carbon cycle and other biogeochemical cycles in the oceans (Kirchman, 2004). Holding an amount of carbon of 660 billion metric tons, oceanic DOM represents the largest reservoir of reactive organic carbon on Earth (Hansell et al., 2009). Given its huge inventory, any change in the rate of DOM mineralization could affect atmospheric CO_2_ concentration with climate consequences; the net oxidation of 1% of the DOM pool would indeed release into the atmosphere an amount of CO_2_ equivalent to the annual anthropogenic emission from fossil fuel combustion and anthropic activities (Le Quéré et al., 2015).

In the open ocean, most of DOM comes from autochthonous production (Jiao et al., 2010; Carlson and Hansell, 2015). In coastal environments, allochthonous inputs from rivers, and runoff may be significant sources of DOM (Hudson and Reynolds, 2007). In addition to natural sources, anthropogenic DOM is directly released into seawater or brought to the coastal areas through atmospheric deposition and freshwater inputs (Pernet-Coudrier et al., 2008; Mostofa et al., 2010). Once in the coastal water, DOM fate depends on biotic and abiotic removal processes. Although abiotic processes, primarily photochemical oxidation (Mopper and Kieber, 2002), may represent an important pathway of DOM removal, the main removal process is related to heterotrophic prokaryotes activity (Azam et al., 1994). DOM is a crucial source of carbon, nitrogen, phosphorus and other essential elements for marine heterotrophic prokaryotes (Azam et al., 1983; Dittmar and Stubbins, 2014; Lønborg et al., 2018). When consumed by heterotrophic prokaryotes, DOM is either used as an energy source through respiration with the production of CO_2_, or converted into bacterial biomass, becoming available for higher trophic levels through the microbial loop (Azam et al., 1983; Cho and Azam, 1990). DOM fate through the microbial loop, i.e. the balance between biomass and CO_2_ production, is determined by various factors.

DOM fate depends on the relationships between HP community, DOM chemical properties and environmental conditions (temperature, salinity). One hypothesis is that DOM intrinsic properties determine its lability and therefore influence its processing by HP communities (Del Giorgio and Cole, 1998; Carlson and Hansell, 2015; Shen and Benner, 2020; Dittmar et al., 2021). Among the environmental conditions, temperature for example controls HP metabolic activity (Pomeroy and Wiebe, 2001). The availability of inorganic nutrients, notably nitrogen and phosphorus, is known for its importance in DOM processing by HP (Del Giorgio and Davies, 2003; Nieto-Cid et al., 2006). Then, biotic characteristics of the marine environment also contribute to DOM fate. First, HP development is under strong top-down pressure by grazers and viruses and strong competition with phytoplankton for inorganic nutrients in marine environments, limiting DOM removal rates (e.g. Lui et al., 2014; Silva et al., 2019). Second, microbial community structure determines its functional and metabolic capacities, therefore influencing DOM processing and fate in a given environment (Pedler et al., 2014; Roller and Schmidt, 2015; Song et al., 2017). Last, DOM reworking by HP communities has been also hypothesized to make DOM more recalcitrant increasing its residence time (persistence) in the oceans (Jiao et al., 2010; Mentges et al., 2019; 2020). All these studies enabled an ecosystemic and dynamic perception of DOM fate, where living and non-living compartments interact, determining DOM persistence in the oceans (Dittmar et al., 2021). To date, factors controlling DOM fate have mainly been considered separately, it is, therefore, necessary to consider these factors together to better predict DOM fate and the resulting consequences on the carbon cycle and the marine ecosystem. In addition, the impact of the numerous natural and anthropogenic inputs on the fate of carbon in coastal areas is not well determined.

The objective of this work is to study the influence of varying chemical properties of DOM and inorganic nutrients on the heterotrophic base of the planktonic food web to provide evidence of the hierarchy of the main drivers controlling DOM fate in coastal marine environments. We adapted a previously-developed transplantation approach (Dignan et al., 2023) to characterize DOM fate from contrasting coastal waters using microbial growth as a proxy. DOM from 2 contrasting environments, with similar dissolved organic carbon (DOC) concentrations but different chemical composition of DOM and inorganic nutrient availability were incubated with two microbial communities presenting contrasted short-term life histories and taxonomic structures. In order to compare the influences of DOM chemical properties, the inorganic nutrients availability and HP community composition, changes in DOM elemental and molecular composition, in the fluorescent fraction of DOM (FDOM), in concentration in DOC, inorganic and organic dissolved nutrient as well as prokaryotic taxonomic diversity were characterized within 22 days.

## 2 Material and methods

### 2.1 Study area

Surface water was collected on May 25^th^, 2021 from two sites with contrasting continental and anthropogenic influences. The civil harbour of Toulon (“T”, 43°7’6.43’’N; 5°56’2.43’’E) was chosen due to its specificities. The coastal water receives discharging water from a large watershed, while having an elevated residence time (Dufresne et al., 2014). The bay is also located at the center of a large urbanized area, hosting leisure and fishing boats. These combined factors provide opportunity to observe exacerbated anthropogenic and continental influences (Coclet et al., 2018). On the other hand, the Niel Bay of Giens (“G”, 43°2’7.12’’N; 6°7’39.03’’E) is less than twenty kilometers away from T, thus exposed to similar climatic and tidal conditions as T. However, G is located on the Giens peninsula, within the Port-Cros National Park. This site is less subjected to continental and human influences because of a smaller watershed, very low anthropogenic activity and continuous stirring by marine currents. These contrasting environmental conditions were expected to demonstrate large differences in DOM pools. Large differences in microbial community structures and functioning have already been reported in the area and are associated with short-term life history (Coclet et al., 2018, 2019, 2020; Dignan et al., 2023).

### 2.2 Sample collection

Seawater was collected using a Van-Dorn sampler and transferred in 20 L LDPE bottles. The materials used for sampling and experimentation have undergone a washing and conditioning protocol before use in order to avoid any contamination (see supplementary material). Water temperature, salinity and chlorophyll a were measured *in situ* using a multiparameter probe (Hydrolab® DS5X). To minimize changes in DOM properties and prokaryotic communities, the samples were brought back in the laboratory within 2 hours and immediately processed in order to launch the experiments as soon as possible.

### 2.3 Experimental design

The incubation experiments consisted in incubating each pool of DOM to the HP community from the same sampling site (Dignan et al., 2023), for a representative evaluation of DOM bioavailability while minimizing manipulations. These experimental conditions were referred to as “TinT” for the harbour community of T in its own dissolved substances and “GinG” for the community of G in its own dissolved substances. Each DOM pool was also incubated with the HP community from the other site in order to evaluate whether the same pool of DOM could experience a different rate (e.g. different bioavailability) with another microbial community. This incubation also allowed us to document community dynamics according to different bottom-up influences. The corresponding experimental conditions were named “TinG” for the community from T in DOM from G and “GinT” for the community from G in DOM from T.

This cross-exposure approach resulted in four experimental conditions, each one carried out in triplicates. For each replicate, 1.8 L of filter-sterile seawater (< 0.2 µm, polyethersulfone filter, Whatman, 47 mm) containing DOM was inoculated with 0.2 L of seawater filtered through a GF/F filter (< 0.7 µm, Whatman, 47 mm) in order to contain the free-living HP community. Incubations were carried out in 2 L fluorinated ethylene propylene (FEP) bottle. All the bottles were incubated at *in-situ* temperature (15°C) and in the dark (to avoid photosynthetic DOM production) for 22 days. To monitor the growth of HP, subsamples were taken immediately after inoculum (T_0_) and after 1, 2, 4, 8, 10, 15 and 22 days of incubation. Aliquots of samples were also sampled at T_0_ and T_22_ to follow dissolved organic carbon (DOC) and nutrients concentration as well as DOM molecular composition, changes in fluorescent DOM (FDOM) and HP community composition. The 2 L bottles were gently mixed before each subsampling, which was carried out taking the necessary precautions to avoid any contamination by ambient microbes and DOM. All subsampling material used has been previously conditioned (See supplementary material).

### 2.4 DOC analyses

Subsamples of 24 mL were filtered through preconditioned 0.2 µm polyethersulfone (PES, 33 mm) syringe filters, collected in preconditioned glass tubes, acidified (10% v/v HCl, analytical grade, Fisher Scientific) and stored at 4°C for DOC concentration measurement. DOC concentrations were determined by high temperature catalytic oxidation using a Shimadzu TOC-VCSN carbon analyser following the method reported by Santinelli et al. (2015). At the beginning and the end of each analytical day, system blanks were measured using Milli-Q water and the reliability of measurements was checked by comparison with a DOC certified reference material (CRM) (CRM Batch #20; nominal concentration of 42.1 ± 1.2 µM; measured concentration 41.6 ± 0.7 µM, n = 112) (Hansell, 2005).

### 2.5 EEM-PARAFAC analyses

Following the same filtration protocol as for DOC, 24 mL of filtered water were stored in preconditioned glass tubes at 4°C for FDOM analyses. The fluorescence excitation-emission matrices (EEMs) were recorded using a Aqualog spectrofluorometer (Horiba) following the method reported by Retelletti Brogi et al. (2020). EEMs are used to distinguish fluorescence signals due to various groups of fluorophores. The excitation wavelength ranged between 220 and 450 nm with a 5 nm increment, the emission was measured between 220 and 600 nm. By using the TreatEEM software (Omanovic et al., 2023), EEMs were elaborated to remove and interpolate Rayleigh and Raman scatter peaks and then normalized to the Raman signal of water, dividing the fluorescence by the integrated Raman band of Milli-Q water measured on the same day of analyses. Fluorescence intensity is expressed in water-equivalent Raman units (R.U.) (Lawaetz and Stedmon, 2009). PARAFAC analyses was performed on all the EEMs collected in the experiments by using MATLAB R2018a (Mathworks, Natick, MA). The final 4-component model was validated by visual inspection of residuals, split-half analysis and the percentage of explained variance (Stedmon and Bro, 2008). The 4 components were characterised as C1_MarHum-like_ (marine humic-like component, Ex/Em: 295nm/400nm), C2_TerHum-like_ (terrestrial humic-like component, Ex/Em: 365nm/465nm), C3_Tyr-like_ (tyrosine protein-like component, Ex/Em: 260nm/312nm), C4_Trp-like_ (tryptophane protein-like component, Ex/Em: 280nm/338nm). C1_MarHum-like_ is similar to humic-like M peak, which is widely observed in coastal water characterised by high microbial activity (Coble, 1996; Stedmon et al., 2011). C2_TerHum-like_ component is considered as a terrestrially derived humic-like products (Coble et al. 2014). C3_Tyr-like_ component is similar to the peak B for tyrosine protein-like sources (Coble, 1996). C4_Trp-like_ component is similar to Tryptophan protein-like T peak (Coble, 1996).

### 2.6 Nutrient concentrations

For nutrient concentration measurements, subsamples (50 mL) were filtered through 0.2 µm PES (33 mm) syringe filters, collected in centrifuge tubes (Falcon) and stored at −20°C. Nitrate, nitrite and phosphate concentrations were measured by colorimetric methods using an automated Technicon Autoanalyser III (Tréguer and Le Corre, 1975) with respective detection limits 0.05, 0.03, 0.02 µmol.L^-1^. Total element concentrations (total nitrogen TN and total phosphorus TP) were determined using the wet-oxidation technique (Raimbault et al., 1999). Dissolved organic nitrogen (DON) and phosphate (DOP) concentrations were empirically determined by subtracting the nitrate and nitrite concentrations from TN and orthophosphate concentrations from TP, respectively.

### 2.7 DOM molecular characterization and data processing

One liter of water sampled at T_22_ was filtered through 0.2 µm pore filter (PES filter, Whatman, 47mm) and collected in 1L FEP bottles. It was then acidified to pH 2 with HCl (32%, puriss p.a. grade). According to the manufacturer’s guidelines, extraction cartridges Bond Elut PPL (Agilent) were activated the day before the extraction by soaking them in methanol (p.a. grade). Immediately before use, the extraction cartridges were rinsed with two fillings of MilliQ 18.2 MΩ water, methanol and acidified MilliQ water (pH 2). The acidied samples were gravity-filtered through the cartridges at flow rates not exceeding 20 mL.min^-1^. The subsample bottles were connected to the cartridges with Teflon tubings (ID 2 mm) and Luer adaptors. The cartridges were subsequently rinsed with at least three cartridge volumes (18 mL) of acidified Milli-Q water for complete removal of salt. The sorbents were dried with ultrapure nitrogen gas for about 5 min, and DOM was immediately eluted with 7 mL of methanol in 24 mL glass tubes. The eluates were stored at 4°C until further analyses (Dittmar et al, 2008; Shen et al., 2016).

DOC concentrations were measured at the inlet and at the outlet of the cartridge to evaluate column recovery (ca. 60% of the total DOC was recovered by the cartridges). The molecular characterization of these extracts was carried out using high-resolution mass spectrometry (HRMS), e.g., Orbitrap Q Exactive (Thermo Fisher Scientific) in infusion mode at the Water Quality Centre (Trent University, Canada). Negative [M − H]^-^ ions were examined at a resolving power of 140,000, using a full MS scan ranging from 100 to 1000. The capillary temperature was set at 320°C and the voltage of sputtering was maintained at 3000 V. The sheath gas pressure was 12 (arbitrary unit). The solution injected (at 10 μL.min^-1^) was diluted to a concentration of 10 mgC.L^-1^ in a mixture of MilliQ 18 MΩ water and methanol (50/50, v/v). Prior to analysis, a routine mass calibration was performed with Pierce™ LTQ ESI Positive/Negative Ion Calibration Solution (88322/88324, ThermoFisher Scientific). Samples were spiked with sodium dodecyl-d25 sulfate (m/z 290.3048) and dioctylsodium sulfosuccinate (m/z 421.2265) as lock masses to ensure ppm-level mass tolerance.

Molecular formulas were assigned using the open access, browser-based software UltraMassExplorer (UME; https://ume.awi.de; Leefmann et al., 2019). The assignments were based on a library of pre-built “02-NOM” molecular formulas. Elementary selection of ^1^H_1-∞_, ^12^C_1-∞_, ^14^N_0-4_, ^16^O_0-∞_, ^32^S_0-1_, ^31^P_0-1_ has been applied. User-defined mass tolerance and mass precision have been set to 1 ppm and four decimal places, respectively. An analytical filter (remove the formula not verified by ^13^C, ^34^S and ^15^N isotopes) and a ratio filter (O/C ≤ 1.2, H/C ≤ 2.5) were also applied. The assigned molecules were classified into eight classes of compounds or 12 elementary combinations. Classes of biochemical compounds are reported as relative abundance values based on C, H, and O counts for the following H:C and O:C ranges: lipids (0 < O:C ≤ 0.3, 1.5 ≤ H:C ≤ 2.5), unsaturated hydrocarbons (0 ≤ O:C ≤ 0.125, 0.8 ≤ H:C < 2.5), proteins (0.3 < O:C ≤ 0.55, 1.5 ≤ H:C ≤ 2.3), amino sugars (0.55 < O:C ≤ 0.7, 1.5 ≤ H:C ≤ 2.2), carbohydrates (0.7 < O:C ≤ 1.5, 1.5 ≤ H:C ≤ 2.5), lignin (0.125 < O:C ≤ 0.65, 0.8 ≤ H:C < 1.5), tannins (0.65 < O:C ≤ 1.1, 0.8 ≤ H:C < 1.5), carbohydrates (0.7 < O:C ≤ 1.2, 1.5 ≤ H:C ≤ 2.5) and condensed hydrocarbons (0 ≤ O:C ≤ 0.95, 0.2 ≤ H:C < 0.8) (Minor et al., 2014). The elementary combinations were CH, CHN, CHNO, CHNOP, CHNOS, CHNOPS, CHNS, CHO, CHOP, CHOS, CHOPS and CHS (Rivas-Ubach et al., 2018). The intensity of a measured compound has been transformed into frequency in order to allow the comparison among the samples.

A molecular lability boundary (MLB) was developed to divide the ultrahigh-resolution mass spectrometry molecular data into two groups encompassing theoretically more or less labile material, derived from calculated hydrogen saturation molecular values (D’Andrilli et al., 2015). DOM constituents above the MLB at H/C ≥ 1.5 correspond to theoretically more labile material in more bioavailable chemical class regions (grouping in protein-, amino sugar-,carbohydrate-, lipid-like regions of the van Krevelen diagram), whereas DOM constituents below the MLB, H/C < 1.5, exhibit theoretically less lability, more recalcitrant character (grouping in more lignin-, tannin-, and condensed aromatic-like regions of the van Krevelen diagram) over all O/C ratios (Minor et al., 2014; D’Andrilli et al., 2015). The percentage of theoretically more labile (MLB_L_) or recalcitrant (MLB_R_) constituents is determined by summing the molecule frequencies categorised with a ratio, respectively, H/C ≥ 1.5 or H/C < 1.5 over all O/C (0.0 – 1.2) multiplied by 100.

### 2.8 Biological analyses

#### 2.8.1 Abundance, flow cytometry

Dedicated 1 mL subsamples were transferred to 1.5 mL Eppendorf tubes, fixed with glutaraldehyde (0.25% final concentration), and stored at −80°C prior to HP enumeration by flow cytometry. After thawing, samples were stained with SYBR Green (1X final concentration) for 15 minutes in the dark and HP were counted with an Accuri C6 flow cytometer (BD Biosciences). Fifty µL were run at a flow rate of 35 µL.min^-1^. Non-fluorescent polystyrene microspheres (Flow Cytometry Size Calibration Kit, Thermo Fisher Scientific) were used as a size standard for daily calibration. Particles considered as HP were smaller than 2.0 µm, exhibited low complexity (low SSC), emitted green fluorescence and no red fluorescence (Grégori et al., 2001). Data were acquired using BD Accuri CFlow Plus software and heterotrophic prokaryotes abundance (HPA) expressed as cells per milliliter (cell.mL^-1^).

#### 2.8.2 Taxonomic diversity, metabarcoding

To study HP community structure, HP were collected by filtering 60 mL subsamples through a 0.2 µm pore filter (Whatman, 47 mm) and the membrane filters were stored at −4°C until further processing. After thawing, they were cut with sterile scissors into small pieces, immersed in 750 µL of sterile saline TSE buffer (50 mM Tris-HCl, 750 mM sucrose, 20 mM EDTA, 40 mM NaCl, pH = 9.0) and horizontally vortexed for 5 min at ambient temperature. Cell lysis was performed with lysozyme, SDS and proteinase K as reported elsewhere (Boström et al., 2004). DNA was precipitated in 70 % ethanol, 0.1 M sodium acetate (pH = 5.2) and linear polyacrylamide (Genelute-LPA, Sigma-Aldrich) (Gaillard et al., 1990). After 2 rinsing steps with 70% ethanol, DNA pellets were air-dried, resuspended in Milli-Q water and stored at − 20°C. Amplification of the V4-V5 region of 16S rRNA gene targeting bacteria and archaea was performed with primers 515F-Y/926R (Parada et al., 2016). Reaction mixtures contained 10 µL of DNA extract, 2X GoTaq Long PCR Master Mix (Promega, Madison, WI, USA) and 0.4 µM of each primer, in a final volume of 60 µL. The PCR program included an initial heating step of 2 minutes at 95°C followed by 30 cycles of 95°C for 30 seconds, 50°C for 45 seconds and 72°C for 45 seconds, and a final extension of 2 minutes at 72°C. PCR amplification efficiency and specificity were checked after migration of 5 µL of PCR products on a 1.5% agarose gel. After purification (Nucleospin Gel and PCR clean-up, Macherey-Nagel), PCR products were paired-end sequenced (2×300 bp) with an Illumina Mi-Seq sequencer by Eurofins (Germany). Sequencing reads were deposited in the National Center for Biotechnology Information Sequence Read Archive (NCBI SRA) under the accession number PRJNA952911. MiSeq raw reads were analysed with *DADA2* (Callahan et al., 2016) in RStudio 2022.7.0.548 (RStudio Team, 2022) under R 4.2.0 environment. Sequences were trimmed according to average Q-scores, then filtered allowing no N, no more than 2 expected errors whatever the read (i.e., forward or reverse), and a minimal Q-score of 3. Taxonomic information was assigned to ASVs with the SILVA v.138 database (Pruesse et al., 2007; Quast et al., 2013). Sequences classified as mitochondria or chloroplasts were removed from the dataset. Extraction and amplification negative controls were sequenced, in case of contamination, the amplified groups were removed from the other samples dataset. One sample (GinT after 4 days of incubation) was also removed from the dataset because it did not have enough sequences.

### 2.9 Statistical analyses

All statistical analyses were performed with RStudio 2022.7.0.548 software (RStudio Team, 2022) under R 4.2.0 environment. All comparisons between initial and final incubation times of each condition were done by Mann Whitney non parametric test of the *vegan* package. All the differences presented as significant in this article had confidence interval of 95%. After verifying the normal distribution and homoscedasticity of the variances with Shapiro and Levene tests, respectively, analyses of variance (ANOVA) were used to compare fluorescence intensity values and HPA values between each incubation time and each condition using *rstatix* package. If the ANOVA detected a significant difference, pairwise comparisons of estimated marginal means with the Bonferroni method for the p-value correction was then used to distinguish which condition differed significantly between the incubation times and between the other conditions.

With ultra-high resolution mass spectrometry data, the fold-change of each condition was calculated by taking the log 2 of the ratio between the data of a condition at the final and initial time. The significance of the temporal variation was assessed using the Mann Whitney non-parametric test of the *vegan* package. The significance of temporal variability versus fold-change of the identified molecules was represented with volcano plots using *dplyr* package (Fig. S.I.1). Molecules showing significant variations were used to construct Van Krevelen diagrams, using *ggplot2* package, representing the molecules either significantly consumed or significantly produced in each experimental condition.

With metabarcoding data, “heat trees” were generated using *metacoder* package (Foster et al., 2017) to pairwise-compare the conditions and identify the taxa most strongly driving the compositional differences. Alpha diversity calculations (Shannon) were performed using *phyloseq* package (McMurdie & Holmes, 2013). For community profiling, beta-diversity based analysis of similarities on Unifrac distances was calculated using the *phyloseq* (McMurdie & Holmes, 2013) and *vegan* (Oksanen et al., 2013) packages. Results were plotted on a principal coordinate analysis (PCoA) graph. Graphical representations were performed using *ggpubr* and *ggplot2* packages.

## 3 Results

### 3.1 Physical-chemical characteristics of the 2 sites

Values for all physical-chemical parameters initially measured at the two sites are provided in Tab. 1 and Tab S.I.1. The *in situ* seawater temperature at G and T was similar and close to 15°C. At T, the salinity was 3 units lower than at G. The chlorophyll-a was five times more concentrated at T than at G (Tab. 1).

**Tab. 1:** Average temperature, salinity, chlorophyll a, DOC, DIN, DIP, DOP and DON concentrations assessed in G and T.

|  | Giens | Toulon |
| --- | --- | --- |
| Temperature (°C) | 15.21 | 14.78 |
| Salinity | 39.9 | 36.9 |
| Chlorophyll a (µg/L) | 0.21 | 1.25 |
| DOC (µM) | 106.4 ± 7.6 | 105.4 ± 2.3 |
| DIN (µM) | 0.27 | 18.77 |
| DIP (µM) | <0.02 | <0.02 |
| DOP (µM) | 0.14 | 0.25 |
| DON (µM) | 6.36 | 5.48 |

### 3.2 Incubation experiment

#### 3.2.1 Nutrients and DOC

DIP values measured initially and during incubations were below the detection limits (< 0.02 µM). DOC concentrations measured at G (106.4 ± 7.6 µM) were similar to those measured at T (105.4 ± 2.3 µM) (Tab. 1). During the incubation, DOC concentrations significantly decreased (p-value < 0.05) in all the conditions to reach similar final values (ranging from 79.8 ± 2.6 µM to 87.8 ± 1.3 µM) (Fig. 1.A). DOP concentrations were initially nearly double at T (0.25 µM) than at G (0.14 µM) (Tab. 1). During incubation, a significant decrease in DOP concentration was recorded in all the conditions (p-value < 0.05) but it was twice stronger in conditions with DOM of T than with DOM of G (Fig. 1.B).

**Fig. 1:**
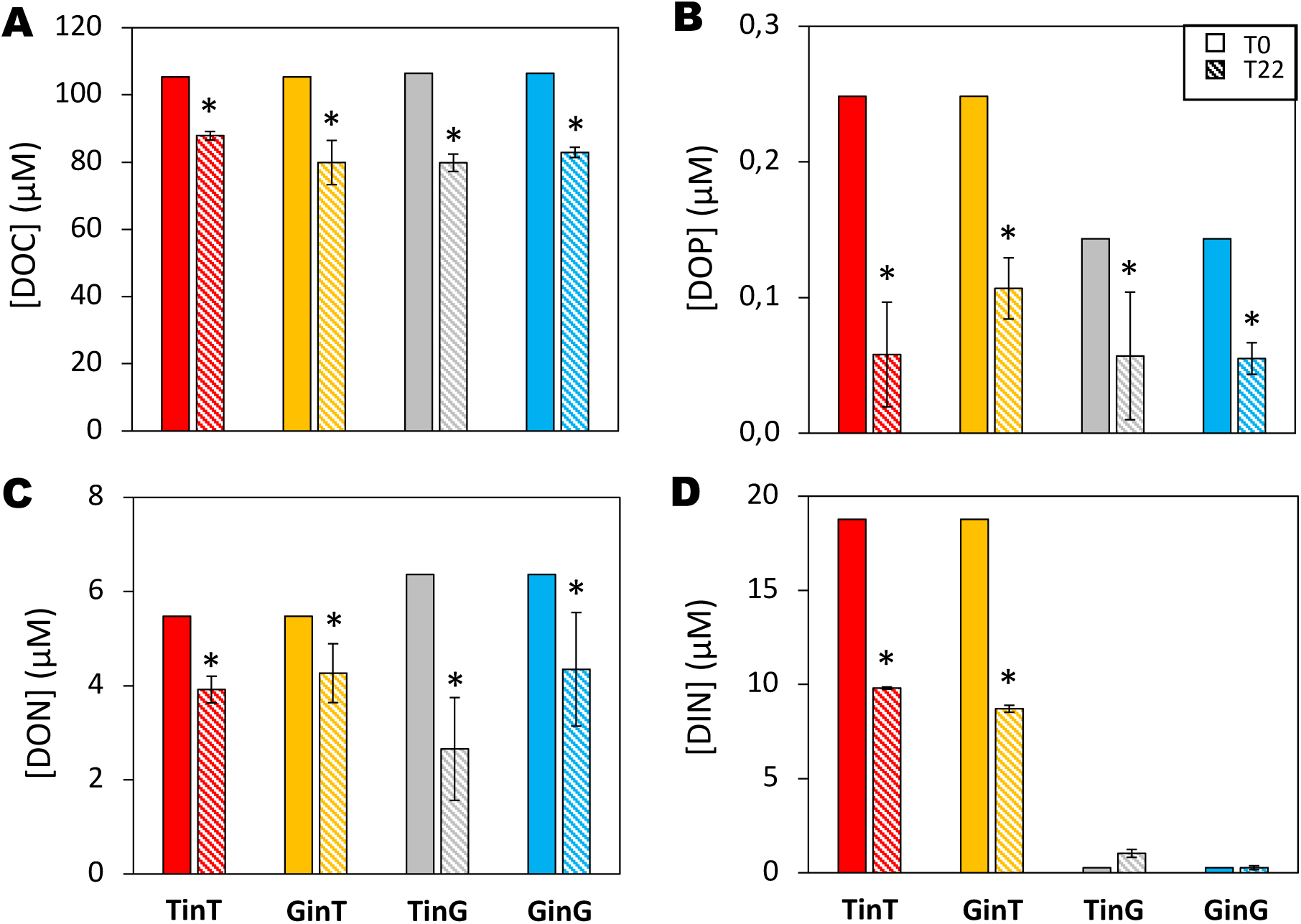
Comparison of abiotic parameters dynamics at the initial (filled colours) and final (fine hatch) time of the incubation experiment for each condition. (A) DOC concentrations, (B) DOP concentrations, (C) DON concentrations, (D) DIN concentrations. Error bars denote standard deviation of triplicated experiments. The stars represent a significant difference between the initial and final incubation time.

DON concentrations measured at G (6.36 µM) were slightly higher to those measured at T (5.48 µM) (Tab. 1). DON concentration significantly decreased in all conditions (p-value < 0.05) but no clear difference between conditions was observable (Fig. 1.C). DIN concentrations were about 70 times more concentrated at T (18.50 µM) than at G (0.26 µM) (Tab. 1). After 22 days of incubation, DIN concentrations significantly decreased (p-value < 0.05) under the conditions with DOM from T, whereas no significant difference was observed under the conditions with DOM from G (Fig. 1.D).

#### 3.2.2 DOM quality

The fluorescence intensity of different components was less than or equal to 0.025 R.U. in G and greater than or equal to 0.039 R.U. in T (Fig. 2). For the conditions TinT and GinT, the fluorescence intensity of C1_MarHum-like_, C3_Tyr-like_ and C4_Trp-like_ significantly decreased (p-value < 0.05) after 22 days of incubation (Fig. 2.B), whereas no significant changes was observed for C2_TerHum-like_ and for all the components in condition TinG and GinG (Fig. 2.B).

**Fig. 2:**
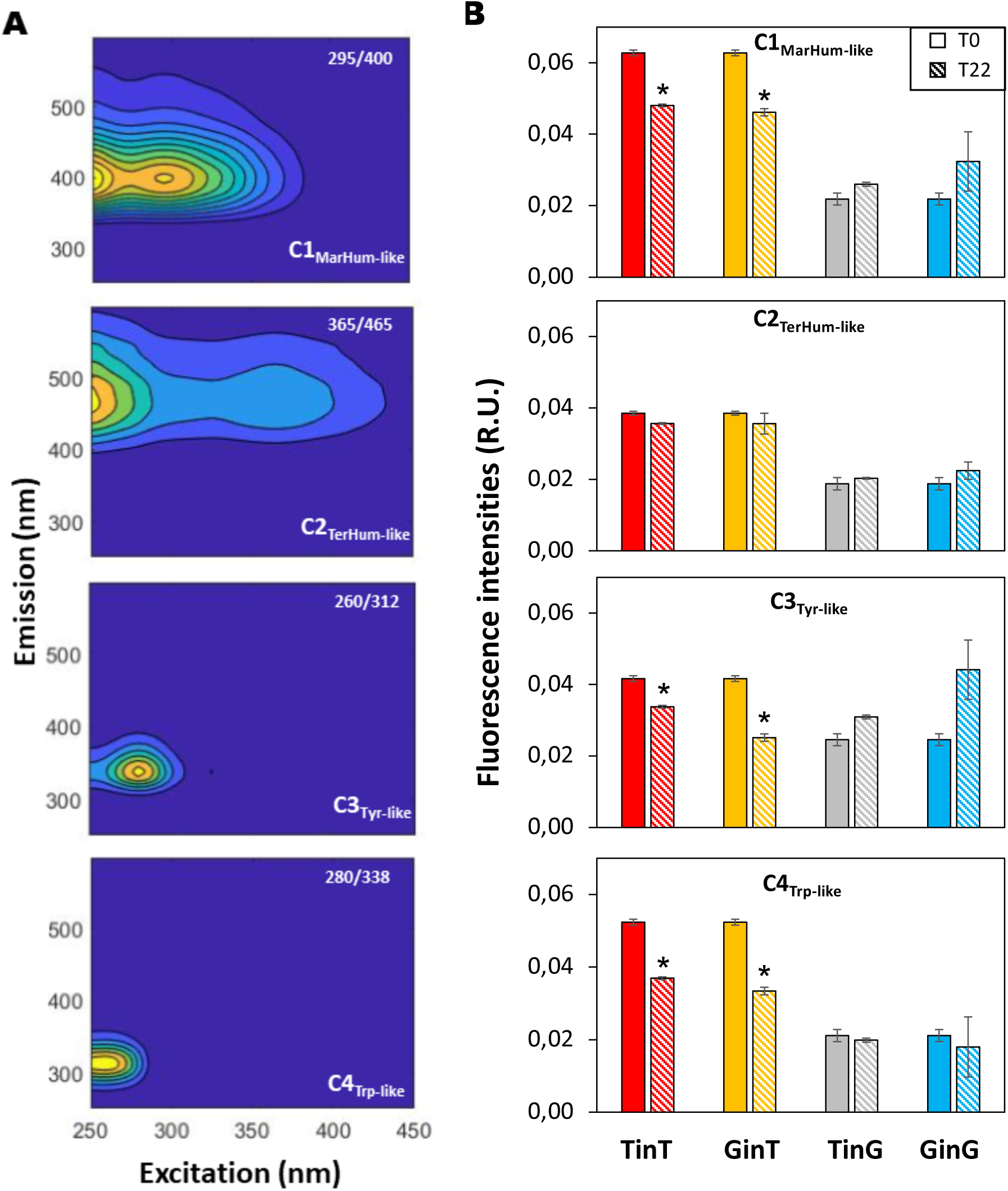
(**A**) Contour plots showing the fluorescence signatures of the four PARAFAC components: C1 marine humic-like, C2 tryptophan-like, C3 terrestrial humic-like, C4 Tyrosine-like. Contour plots of components C1 - C4 are ordered by decreasing percent explained, with emission wavelength on the y-axis, excitation wavelength on the x-axis, and shading representing the relative intensity of emission. (**B**) Changes in fluorescence intensity between the beginning (filled bars) and the end (hatched bars) of the incubation for each condition. The stars indicate a significant difference with initial incubation time.

HRMS analysis identified 5,551 molecules in the different samples. PCoA strongly discriminated DOM molecular composition of T and G, both at the beginning and the end of the experiment. In the conditions with DOM from T, DOM molecular composition of the initial and final incubation times were also well discriminated in the PCoA (Fig. S.I.2.A). According to the H:C ratio of each molecular formula, both sites contained more theoretically labile molecules (≥ 69%) than recalcitrant ones (Fig. S.I.3). Initial DOM from T and G was mainly composed of proteins, carbohydrates and amino sugars. However, the differencial analysis between the two initial DOM pools showed that the initial DOM from T was enriched in proteic compounds, carbohydrates and amino sugars with respect to DOM from G. DOM from G was enriched in lignin-like compounds compared to T (Fig. S.I.4).

Toward the end of the incubation, only 39 to 266 molecules showed a significant decrease in their relative intensity, suggesting the consumption of these molecules by heterotrophic prokaryotes. The proportion of significantly consumed molecules was higher with DOM from T (19.8 and 18.9% in TinT and GinT conditions, respectively) than with DOM from G (9.7 and 1.8% in TinG and GinG conditions, respectively). Among the significantly consumed molecules, more than 77% corresponded to theoretically labile molecules in conditions with the DOM of T while in conditions with the DOM of G, only 33 to 53% corresponded to theoretically labile molecules. Among the significantly consumed molecules, in all the conditions, 90% contained 2 to 4 nitrogen atoms, 60.1 ± 4.7% contained a phosphorus atom and approximately 50% contained a sulfur atom (Fig.S.I.5). The two heteroatomic molecular groups of species containing oxygen, nitrogen and sulfur (CHNOS) or oxygen, nitrogen and phosphorus (CHNOP) represented 80% of the significantly consumed molecules in all conditions except GinG, where other groups such as CHNOPS, CHNO and CHOPS were preferentially consumed (33% and 20% respectively) (Fig.3).

**Fig. 3:**
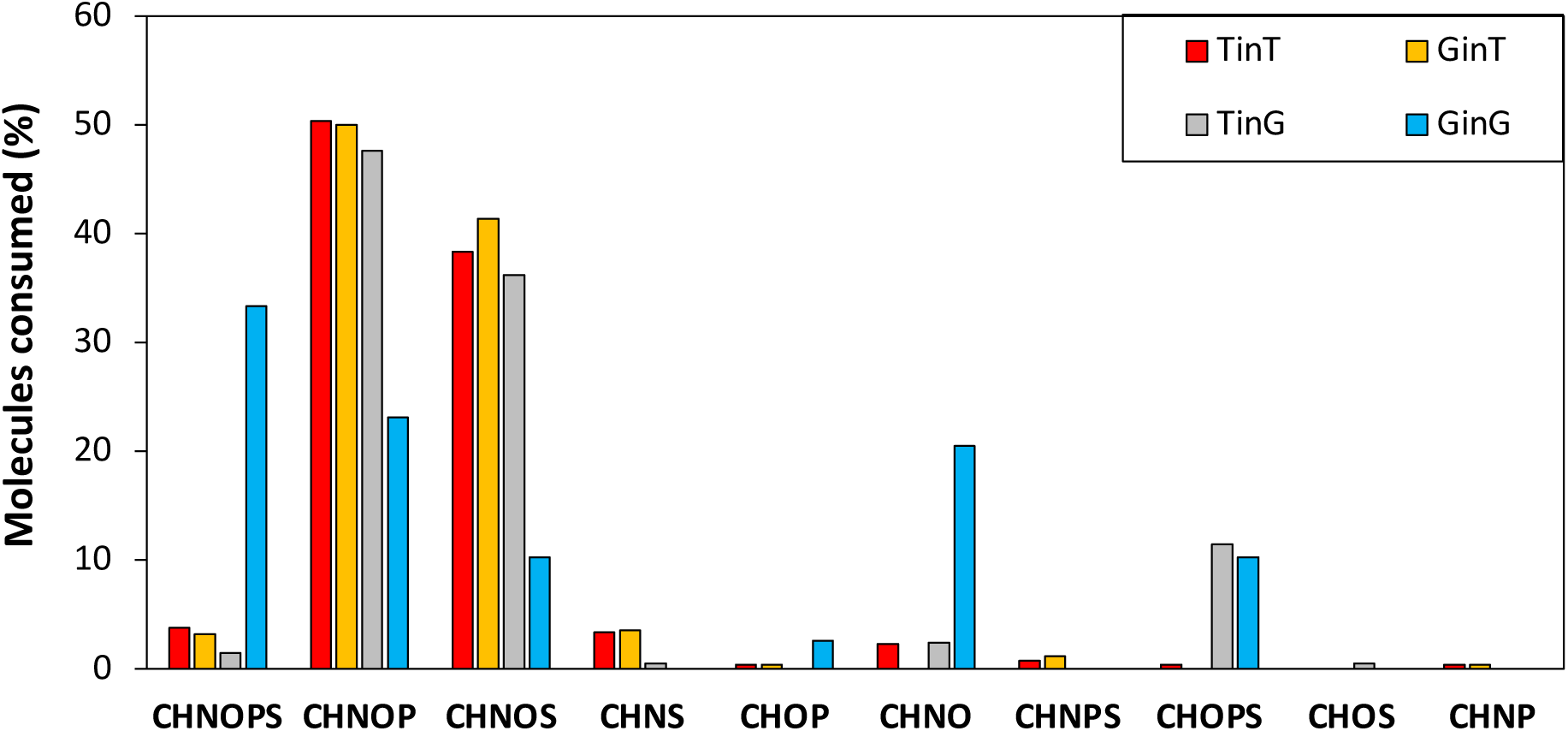
Proportion of each elementary formula among the DOM molecules consumed in each condition. The different heteroatomic molecular group can contain atoms of carbon (C), hydrogen (H), nitrogen (N), oxygen (O), phosphorus (P), and sulfur (S).

The preferentially consumed molecules mostly corresponded to those enriched in either T or G. Indeed, with DOM from T, the most consumed molecular groups were carbohydrates (34.6 ± 3.3% of molecules consumed), proteins (18.2 ± 2.9%) and amino sugars (15.2 ± 3.3%) (Fig. 4). With DOM from G, the preferentially consumed molecules belonged to the molecular groups of lignin (35.6 ± 11.2%) and unsaturated hydrocarbons (19.1 ± 2.0%). It is noteworthy that a significant consumption of proteins (2.8%) and amino sugars (12.8%) was observed in TinG condition but not in GinG (Fig. 4).

**Fig. 4:**
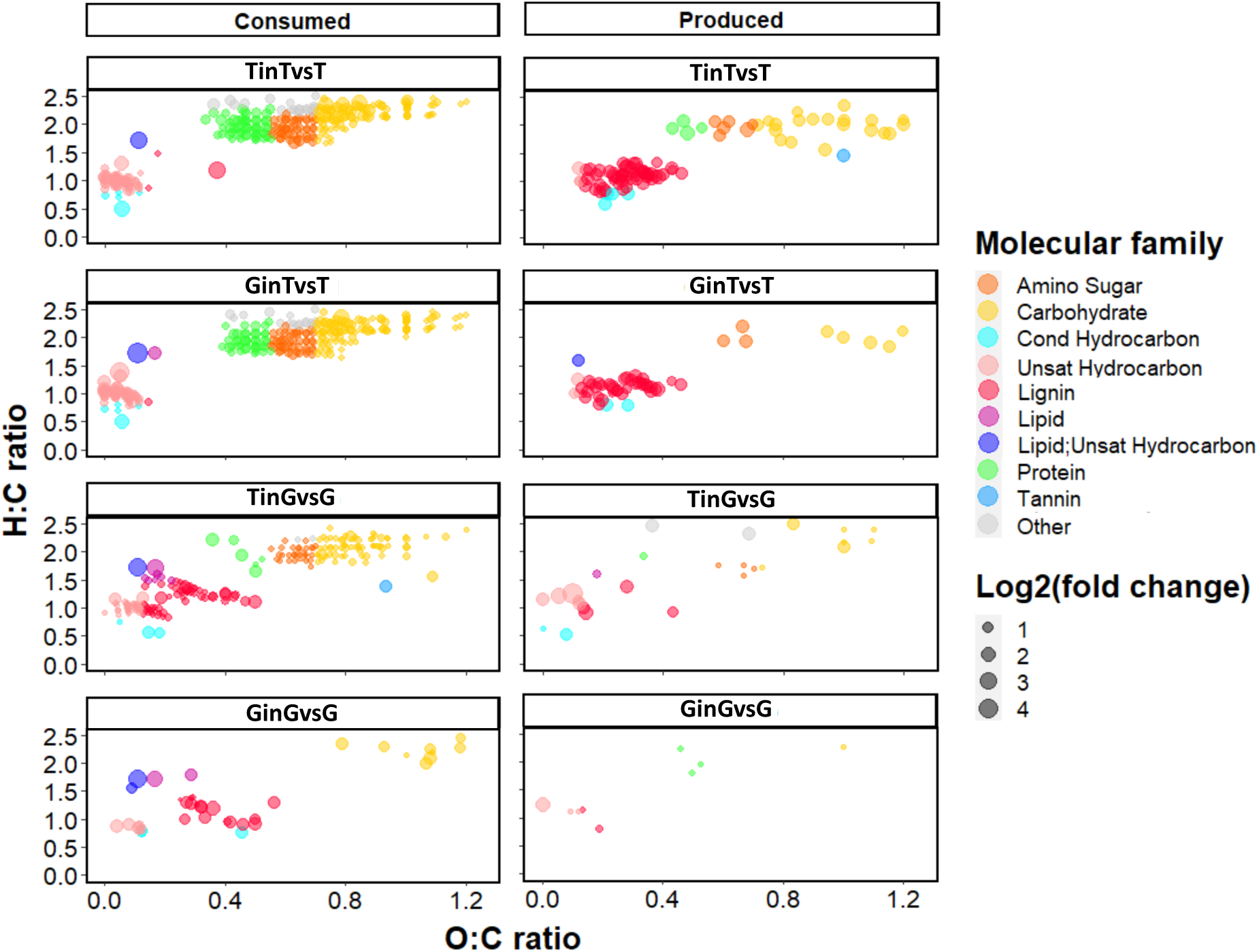
Van Krevelen diagram showing the dynamics of the molecules composing dissolved organic matter under each condition. The left panels represent the molecules significantly consumed during the incubation while the right panels represent the molecules significantly produced during the incubation. The molecules are coloured according to their molecular family. The size of the dots is proportional to the variation factor of a molecule between the initial and final incubation time.

In contrast, we observed a significant enrichment of some molecular groups during the experiment. With DOM from T, the most produced molecular groups were carbohydrates (16.0 ± 8.7%) and lignin (65.1 ± 10.4%) (Fig. 4 and Fig. S.I.6). It is noteworthy that these compounds were the most enriched ones in the other initial DOM pool. With DOM from G, the most produced molecular groups were unsaturated hydrocarbons (26.3 ± 9.9%) and lignin (18.8 ± 4.8%) (Fig. 4 and Fig. S.I.6).

### 3.4. Microbial community

During the incubations, HPA increased in all conditions. Several growth phases were observed: an exponential growth within the 2-4 days of incubation, then the growth slowed down and the abundance remained stable indicating that the community reached the carrying capacity of the system (Fig. 5). DOM from T supported a five-times larger HP carrying capacity than DOM from G, no matter the origin of HP.

**Fig. 5:**
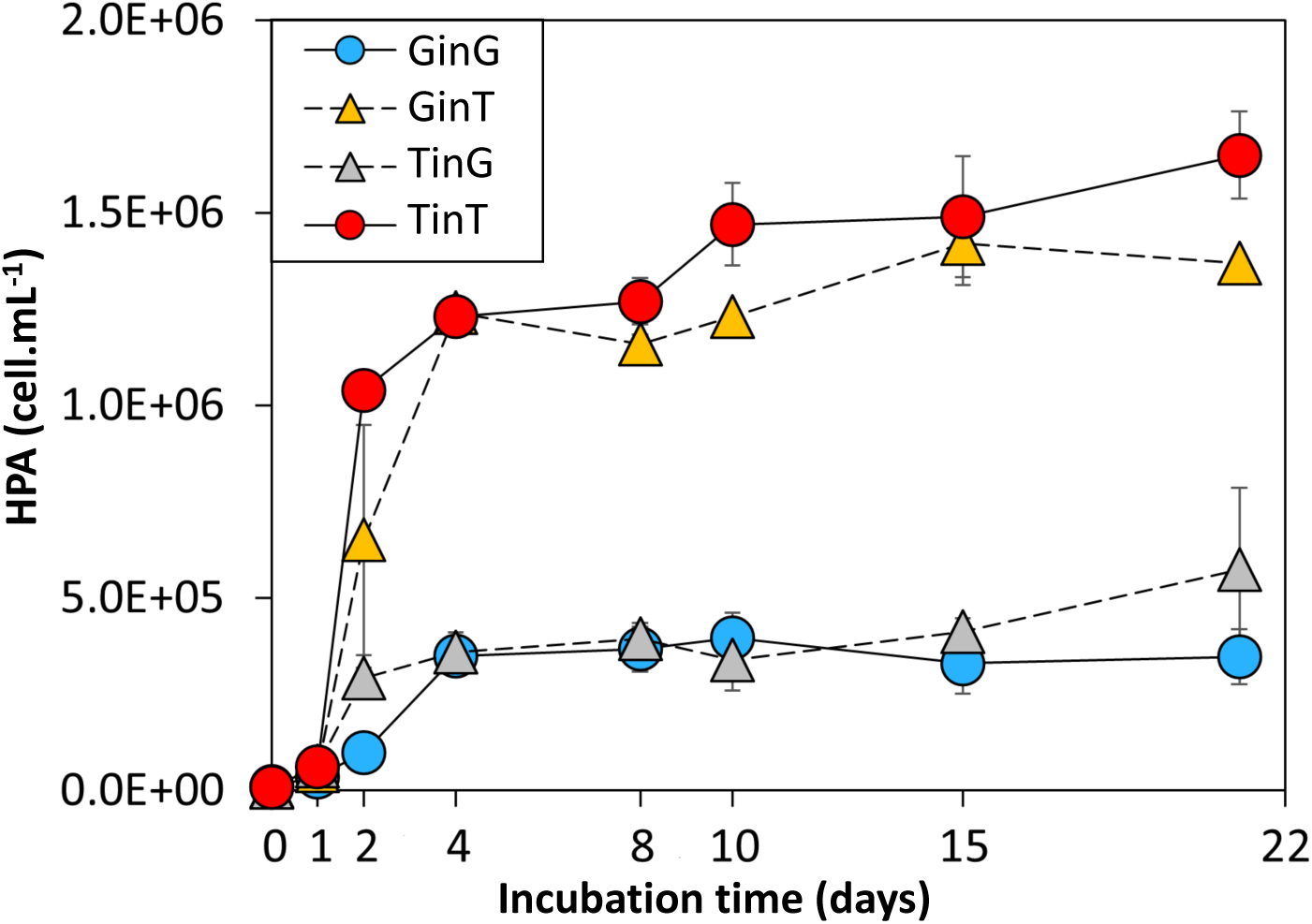
Heterotrophic prokaryotes abundance (HPA) for each condition during the incubation experiment. Errors bars refer to the standard deviation among the 3 replicates.

Initially, the Shannon index was significantly higher for the community from G than for the one from T. In conditions with DOM from T, the Shannon index remained unchanged during the incubation, whereas it significantly decreased with DOM from G (Fig. 6.A). Corresponding partly to alpha diversity results, unifrac distance analysis strongly discriminated initial HP community structures (Fig. S.I.4.B). It also discriminated well the initial (T0) and final (T22) incubation times. It revealed a notable discrimination of the HP community structure at the end of the experiment: unlike the GinG condition, community structure in the condition TinG was not significantly different from conditions with DOM from T.

**Fig. 6:**
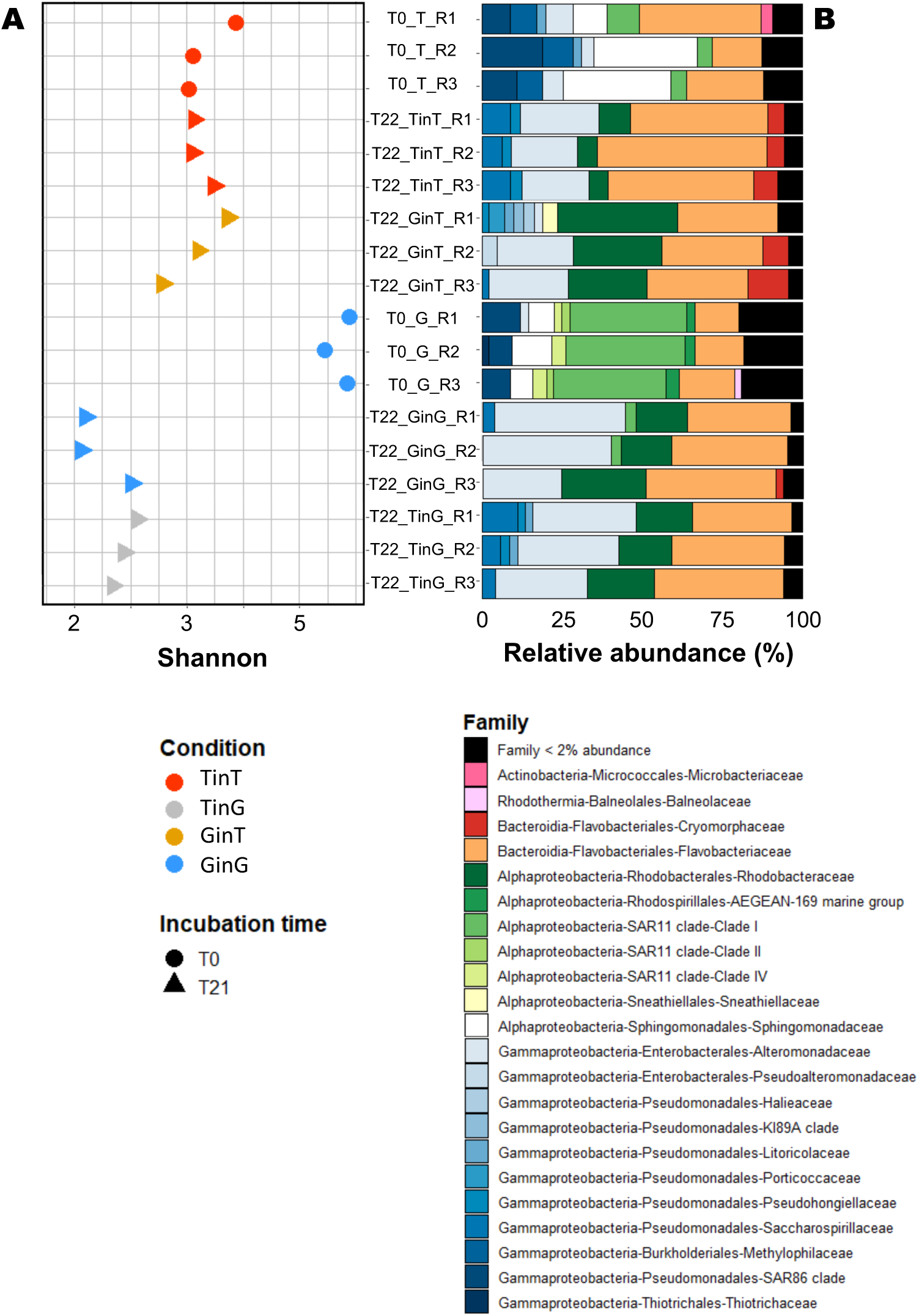
**(A)** Shannon index values calculated from 16SrRNA OTU table for each condition at initial incubation time (circle), after four days of incubation (square) and at final incubation time (triangle). **(B)** Structure of procaryotic community at the family level for each condition. Each bar represents the relative abundance of a family in the studied samples. Families never representing more than 2% of the community were gathered under the black bar.

The most abundant prokaryotic phyla were Proteobacteria (50.6% of total reads) and Bacteroidota (17.9% of total reads). Archaea were poorly represented in the dataset (< 2% in all samples). Initially, prokaryotic communities from G were dominated by families of the SAR11 Clade 1 (Alphaproteobacteria, 36.0%) followed by Flavobacteriaceae (Bacteroidota, 15.4%), while prokaryotic communities from T were dominated by Flavobacteriaceae (25.7%), Sphingomonadacae (Alphaproteobacteria, 25.4%) and families from the SAR86 clade (Gammaproteobacteria, 12.8%) (Fig. 6.B). After 22 days of incubation, the relative abundance of Alteromonadaceae (Gammaproteobacteria) sharply increased from 1-6% to 45-60% and dominated in all conditions. Considering that community structure mainly differed between GinG and all other conditions at the end of the experiment, a heat tree showing differential prokaryotic enrichments at all taxonomic levels from the phylum to the genus between GinG and TinT was computed (Fig. 7). This analysis emphasized the development of Pseudomonadales (Pseudohongiellaceae and Saccharospirillaceae) and Flavobacteriales (Flavobacteriaceae) with DOM from T that promoted the highest HP growth. The relative abundance of Flavobacteriaceae was maintained after 22 days of incubation (9 to 20%) with DOM from T, making this family the second most abundant one after Alteromonadaceae (Fig. 6.B). In contrast, DOM from G promoted the development of Rhodobacterales (Rhodobacteraceae), especially *Nereida ignava*, whose abundance was initially <1% (Fig. 6.B). Some Enterobacterales (Alteromonadaceae) also appeared favoured while the relative decrease of the initially abundant clade SAR11 appeared limited when compared to TinT (Fig. 7).

**Fig. 7:**
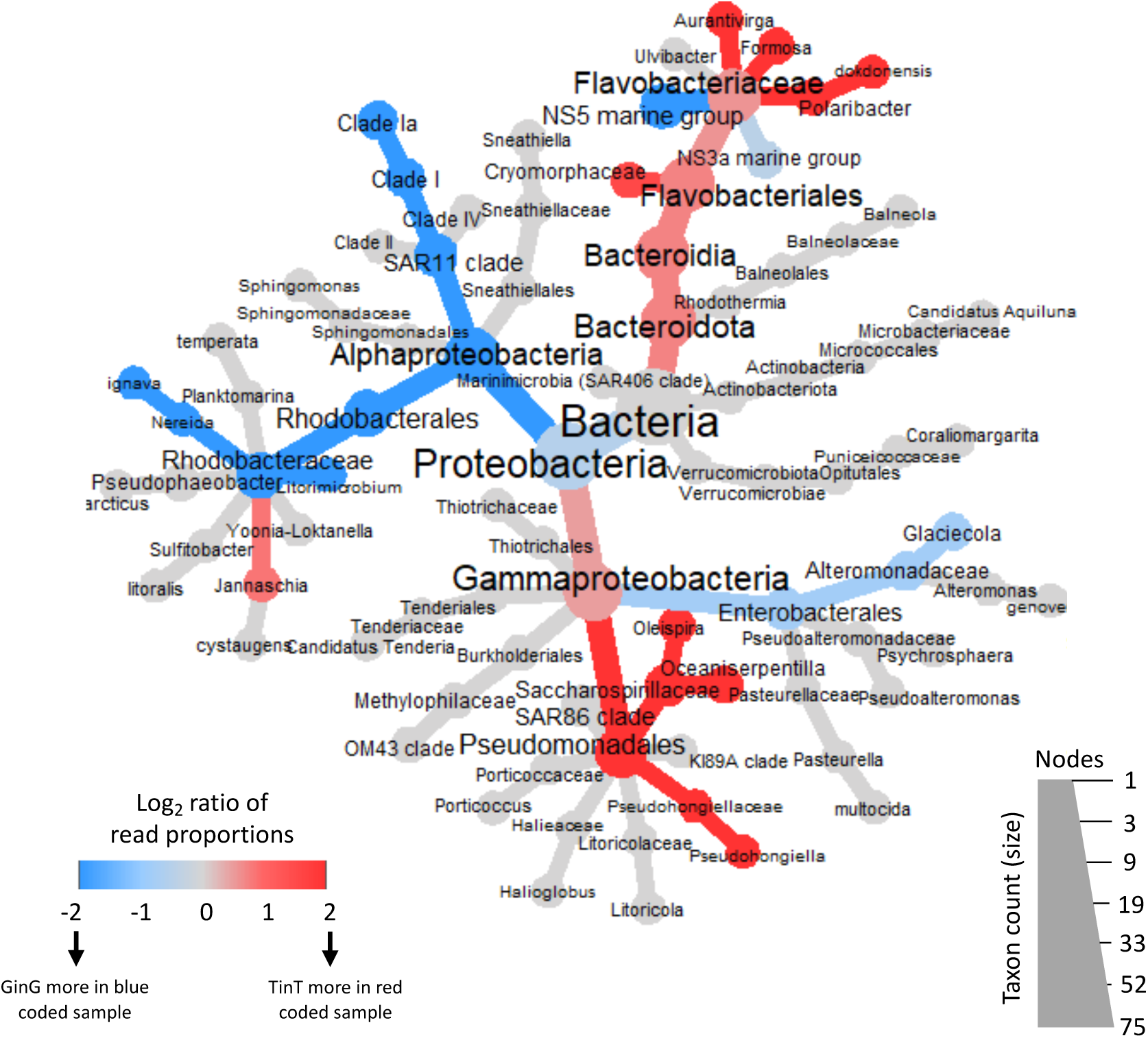
Heat tree illustrating the overall taxonomy of the heterotrophic procaryote community of GinG condition relative to that of TinT condition. Only taxa with a relative abundance greater than 1% were considered in the construction of the heat tree. Nodes represent taxonomic levels and branches represent the affiliations. Color changes represent the difference in log2 ratio of median proportions of reads between GinG and TinT conditions. The blue nodes are more represented in the GinG condition, while red nodes are more represented in the TinT condition. The gray nodes appear equally in both conditions.

## 1 Discussion

### 4.1. Similar DOM lability but different DOM fate

Even if different hypotheses have been formulated, a clear understanding of the hierarchy of quantitative and qualitative DOM properties in controlling DOM fate in oceans is still missing (Shen and Benner, 2020; Lennartz and Dittmar, 2022; Shen and Benner, 2022). In this study, the average DOC concentrations at the two sampled sites were similar depiste differential influence from the continents and anthropogenic activities; this situation offers a unique opportunity to hierarchize the main drivers controlling DOM fate by freeing ourselves from variations in DOC concentration. DOC concentrations were by the way in line with values commonly found in surface coastal waters of the Mediterranean Sea and more specifically in the studied area (Santinelli, 2015; Coclet et al., 2019; Dignan et al., 2023). DOC removal was quite low (∼ 10 % in all the experimental conditions) suggesting a similar percentage of potential labile DOM in the two pools (G and T) on the short (22 days) temporal scale. The removed fraction (10 % of the initial pool) is similar to what is observed in a variety of marine environments and with different initial DOC concentrations (for more information, see Retelletti-Brogi et al. 2021 and references herein) and suggests an intriguing quantitative uptake of DOM that would go beyond concentration-driven control (Lennartz and Dittmar, 2022). Furthermore, in our experiments, DOC removal did not appear to depend on DOM composition, thus contradicting previous work (Del Giorgio and Cole, 1998; Kowalczuk et al., 2013; Shen and Benner, 2020). Similarly, DOC removal did not seem to depend on inorganic nutrient availability or community structure, contradicting again the literature (Del Giorgio and Davies, 2003; Nieto-Cid et al., 2006; Carlson and Hansell, 2015). Thus, with strongly contrasted qualitative DOM properties but similar DOC removal, our results tend to support a weak control of DOM properties on DOM lability.

Measuring DOM lability relies on our ability to precisely detect small variations in DOC concentration. Alternative methods have been proposed in the literature, such as the use of H:C ratio of the molecules as a proxy of their individual lability (D’Andrilli et al., 2015). In our study, the molecular characterization carried out by HRMS showed that both DOM pools were composed of a wide variety of molecules, more than 57% of these molecules were predicted to be labile according to the molecular lability boundary (MLB). These so-called labile molecules were mainly distributed into two groups (CHNOP and CHNOS). However, under the experimental conditions of our study, microorganisms consumed only 1.8 to 19.8 % of the total molecules. This result is in agreement with the low fraction of DOC removed during incubations (∼10%). On the other hand, among the consumed molecules, 23 to 67% were defined as non-labile according to the MLB. This observation highlights the gap between prediction of molecular lability based on molecular formulae and the observed consumption of compounds by natural marine microbial communities and once again stresses the high complexity behind DOM persistence in the oceans.

### 4.2. N and P supplies influence DOM fate under equivalent DOC concentration

It is notheworthy that despite similar percentage of potentially labile DOC in two DOM pools, we observed large differences in HP growth, suggesting that the 2 communities used DOM for different purposes. Heterotrophic procaryotes can perform three major functions when processing organic matter: (1) production of new bacterial biomass (bacterial production), (2) respiration (bacterial respiration), and/or (3) DOM reworking (Jiao et al., 2010). In this work, for the same quantity of carbon consumed, the growth of the communities was different depending on the origin of DOM. It suggests different bacterial growth efficiencies and relatively higher use of DOM from T for biomass production, whereas DOM from G was mainly used for bacterial respiration.

This different DOM fate can be explained by the different quality of DOM and/or the different initial concentrations and removal rates of DIN and DOP in T and G. Indeed, although being in line with previous observations in the area, the observed variations spanned from typical Mediterranean off shore values to the highest range of concentrations observed in Toulon Bay (Bogé et al., 2012; Coclet et al., 2019). Where these nutrients were the most concentrated, they were also the most consumed. The supply of macronutrients such as nitrogen and phosphorus had previously been demonstrated to have a major influence on DOM fate (Del Giorgio and Davies, 2003; Nieto-Cid et al., 2006). Indeed, microorganisms regulate the balance between anabolism and catabolism of organic substrates according to their stoichiometric needs.

In the Mediterranean Sea, nitrogen and phosphorus are known to be limiting factors in DOM recycling processes (Pinhassi et al., 2006; Bogé et al., 2012). It is noteworthy that the studied water was highly depleted in phosphate, without any detectable enrichment in the water from the harbour where DIN was strongly enriched. Thus, in our work, DOP represented the main P supply, as demonstrated to a lower extent in previous work in Toulon Bay or in the Gulf of Lion (De Madron et al., 2003; Bogé et al. 2012). With the consumption observed in parallel to HP growth, our study demonstrated the importance of DOP for HP growth in Mediterranean waters, even in more nutritive harbour waters.

Considering that we observed the same result, i.e. no difference in DOM lability but different DOM fate, with two different microbial communities, we hypothesized that the highest availability of DIN and DOP is the main driving factor of DOM fate in phosphate-depleted North-Western Mediterranean coastal waters showing similar DOC concentrations. However, higher availability in DIN and DOP does not seem to trigger DOM lability.

### 4.3. Insights into the influence of DOM molecular composition on its fate

An accurate evaluation of DOM molecular diversity could provide new insights on its control on DOM fate. As a first approach, we identified the main fluorescent components in the DOM pools and studied their dynamics during the incubations. Marine humic-like and protein-like substances showed high initial fluorescence intensities and appeared more consumed in T where the highest HP growth was also observed. Protein-like compounds, as well as their degradation products, are considered a biodegradable fraction of DOM (Bachi et al., 2023). Our observations are in line with these papers, since fluorescence of protein-like compounds significantly decreased in TinT and GinT, suggesting the presence of labile DOC that can be readily assimilated by the microbial communities (Fellman et al., 2009a, b). Humic-like compounds are usually divided into (i) terrestrial humic-like often associated with decomposed plant material and soil of terrigenous origin (McKnight and Aiken, 1998) and (ii) marine humic-like associated with complex and degraded DOM linked to autochthonous biological activity in the water column (Romera-Castillo et al., 2010). It is now recognized that both terrestrial and marine humic-like substances can be directly released by phytoplankton (Bachi et al., 2023). In our experiment, the higher fluorescence and consumption of these components in the water from T than from G supports an influence of FDOM quality on DOM fate.

HRMS data allowed us to evidence the molecular families preferentially removed and/or released by microbial communities during their growth. A significant enrichment in proteins, amino sugars and carbohydrates families was observed in T, corresponding to the DOM pool sustaining higher HP growth. This observation is in agreement with previous studies showing that proteins, amino sugars and carbohydrates in dissolved free forms can be readily assimilated by bacteria and metabolized at high growth efficiencies (del Giorgio and Cole, 1998; Rosenstock and Simon, 2001). However, we also observed that a large number of molecules belonging to these molecular families and initially present in both DOM pools were not consumed during the incubations. Thus we assume that belonging to these molecular families is not a sufficient proof of their systematic use by microbial communities which present clear preferences for specific compounds. Only a part of the molecules belonging to these families are metabolized and contribute to the HP growth capacities and DOM fate. Similarly, numerous proteins, aminosugars and hydrocarbons were present in the DOM pool of G, but were poorly consumed. With this DOM pool, microbial communities consummed preferentially lignin-like compounds. These compounds are mainly considered as refractory molecules of terrestrial origin, although they can be produced by phytoplankton (Zangrando et al., 2019) and *Posidonia* (Kaal et al., 2018). Our experiment suggest that in particular conditions, lignin-like compounds could be preferentially used by marine planktonic HP communities, instead of thought-to-be more available subtrates.

The consumption of certain molecules present in DOM from T resulted in the significant production of various metabolites. It has been demonstrated that heterotrophic prokaryotes can rapidly remove simple compounds and release DOM with a more complex chemical composition (Ogawa et al., 2001). In this study, microorganisms indeed converted amino sugars and proteins into more oxygenated and less hydrated (more unsaturated) structures. Interestingly, these produced compounds belong to the molecular families that were initially enriched in the DOM pool from G site but poorly consumed and resulting in low growth during our experiment. These results suggest that microbial community metabolism can modify the molecular structure of a given DOM pool, therefore influencing its fate through the microbial loop but not necessarily modifying its lability. Such microbial reworking could correspond to the first steps on the way to the production of recalcitrant DOM (Jiao et al., 2010)

### 4.4. HP dynamics appears to respond to DOM changes more than to control DOM fate, under the studied experimental conditions

The diversity and initial composition of microbial communities were different between the two sites, as expected from previous reports (Coclet et al. 2019, 2020). However, these contrasts did not affect DOM removal rates and bacterial growth efficiencies. Community structure was hypothesized to represent a potential driver of DOM fate in marine environments since different microbial taxa could harbour different genetic potential and express different enzymatic abilities (Pedler et al., 2014; Roller and Schmidt, 2015; Song et al., 2017).

Our work showed that, on the contrary, HP communities were controlled by dissolved resources under our experimental conditions. During incubations, alpha diversity quickly decreased in all conditions and the community structure was deeply modified. We are aware that the dilution of HP performed in these marine cultures can deeply alter community structure and functioning through bottle effect (Cram et al., 2015). The experimental conditions allowed the development of several groups and in particular of Alteromonadaceae. Members of this family are known as opportunists having diverse genomic and metabolic potentials, commonly making them the most dominant group especially in conditions with a sufficient supply of labile organic material such as that found during phytoplankton bloom conditions (Xu et al., 2022). Alteromonadaceae are *r*-strategists that can quickly make use of available substrates (Pedler et al., 2014). The experimental conditions chosen in this study allowed to reduce competition and predation and thus favoured the equivalent development of these opportunistic species in all conditions, contributing to the subsequent drop in alpha diversity.

Besides the short-term massive development of Alteromonadaceae in all conditions, long-term differentiation of community structure was observed. While supporting higher growth, dissolved substances from T promoted the development of Gammaproteobacteria, especially Pseudomonadales and some specific Bacteroidota from the Flavobacteriaceae family. Bacteroidetes are known to be abundant when fresh DOM is produced in large quantities, such as during and after algal blooms (Pinhassi et al., 2004), showing a preference for the consumption of polymers rather than monomers (Cottrell and Kirchman, 2000). The Flavobacteria are particularly specialized in the degradation of proteins (Fernández-Gómez et al., 2013). Their enrichment in incubations with DOM from T was therefore consistent with the consumption of protein-like substances. The dominance of the Bacteroidetes group is often associated with a high nutrient concentration and high bacterial production (Kirchman, 2002), which is in agreement with our observations. Gammaproteobacteria can be selected by high concentrations of labile DOM (Elifantz et al., 2007). This group is known to mainly contribute to the assimilation of sugars (Haider et al., 2023) which could explain the high consumption of amino sugar and carbohydrates in the conditions with DOM from T.

Conversely, supporting lower HP growth, water from G favored the maintenance of typical and initially predominant Alphaproteobacterial oligotrophs such as the SAR11 clade, known to thrive under less nutritive conditions (Mehrshad et al., 2018). This clade is known to mainly use DOC to produce energy (respiration) rather than for biomass production (Sun et al., 2011), and could thus have contributed to the observed low bacterial growth. Then, the assimilable substances from G allowed the development of Rhodobacterales from the Rhodobacteraceae family. More precisely, we observed a specific selection of *Nereida ignava*, a chemoheterotrophic representative from the Roseabacter group known to be poorly reactive to the most used substrates in minimal culture media (Pujalte et al., 2005). The Roseabacter group is well known for its tremendous metabolic diversity (Daniel et al., 2018), and some of its members have recently been suspected of tackling semi-labile DOM (González-Sánchez et al. 2024). More generally, the dominance of Rhodobacteraceae in seawater is often associated with the presence of more complex substrates requiring specific transporters and enzymes for their degradation (Sun et al., 2011; Hernandez et al., 2021). On those bases, we hypothesize that Rhodobacteraceae, especially members from the Roseobacter clade such as *Nereida ignava,* could have been selected by the scarcity of DOM yielding important HP growth. Such selection could have contributed to the degradation of rather recalcitrant organic compounds of the family of lignins.

## Acknowledgments

This project was financially supported by the European Interreg Italy-France Maritime 2014– 2020 Project «GEREMIA» (Gestione dei reflui per il miglioramento delle acque portuali), and by Toulon University, the CARTT of the University Institute of Technology of Toulon and Toulon Provence Mediterrannée through project “C-OMICS” (Coupling OMICS approaches for in-depth study of human-induced responses of microbial communities). Clara Dignan received a Ph.D. fel-lowship from the French Ministère de l’Enseignement Supérieur et de la Recherche and this paper is a part of her Ph.D.

