## Supplementary material for "Unveiling the control of N and P on DOM fate in a Mediterranean coastal environment"

### Pre-cleaning protocol

The water samples were taken with a horizontal van Dorn type water sampler (Wilco, model Beta) previously cleaned with acid (10% v/v HCl, analytical grade, Fisher Scientific) and thoroughly rinsed with Milli-Q water (18.2 MΩ, Millipore) in the laboratory. In the field, the sampler was rinsed with seawater from the site to be sampled before sampling.

Fifteen liters of seawater is stored in a twenty-litter bottle and four litter is stored in a fluorinated ethylene propylene (FEP) bottle. These containers are, in the laboratory, previously rinsed three times with Milli-Q water (18.2 MΩ, Millipore), sterilized with acid (10% v/v HCl, analytical grade, Fisher Scientific) for 24 hours under stirring , rinsed again three times with Milli-Q water (18.2 MΩ, Millipore) then filled with acid (0.1% v/v HCl, Trace Metal Grade, Fluka). In the field, the FEP container and bottle were rinsed three times with seawater from the site to be sampled.

All the FEP bottles used followed the same conditioning protocol described above.

Dissolved substances and heterotrophic communities were isolated by filtration during the experiment. In order to avoid possible contamination of the carbon filtrates, the 0.2 µm polyesthersulfone filters (Whatman, 47 mm) were previously washed with 100 mL of acid (10% v/v HCl, analytical grade, Fisher Scientific), rinsed with 1L of Milli-Q water (18.2 MΩ, Millipore), then packaged with 150 mL of the sample to be filtered. The GF/F glass fiber filters were previously calcined (450°C, 6h), rinsed with 1L of Milli-Q water (18.2 MΩ, Millipore) then conditioned with 150 mL of the sample to be filtered.

The 24 mL glass tubes used for the DOC concentration, DOM fluorescence intensity analysis and DOM molecular characterization were previously rinsed three times with Milli-Q water

(18.2 MΩ, Millipore), sterilized with acid (10% v/v HCl, analytical grade, Fisher Scientific) for 24 hours, rinsed again three times with Milli-Q water (18.2 MΩ, Millipore) then calcined (450°C, 6 hours). At the time of sampling, they were rinsed three times with the sample before being filled.

**Tab. S.I.1: Average (± SD) intensity of DOM fluorescent compounds, number of molecules qualifies as labile ($MLB_L$) and recalcitrant ($MLB_R$) assessed in G and T.**

| | Giens | Toulon |
|---|---|---|
| $C1_{MarHum\text{-}like}$ (R.U.) | 0.022 ± 0.002 | 0.063 ± 0.001 |
| $C2_{TerHum\text{-}like}$ (R.U.) | 0.019 ± 0.002 | 0.039 ± 0.001 |
| $C3_{Tyr\text{-}like}$ (R.U.) | 0.025 ± 0.001 | 0.042 ± 0.001 |
| $C4_{Trp\text{-}like}$ (R.U.) | 0.021 ± 0.001 | 0.052 ± 0.001 |
| $MLB_L$ (%) | 51.72 ± 0.69 | 57.49 ± 3.29 |
| $MLB_R$ (%) | 48.27 ± 0.70 | 42.51 ± 3.29 |

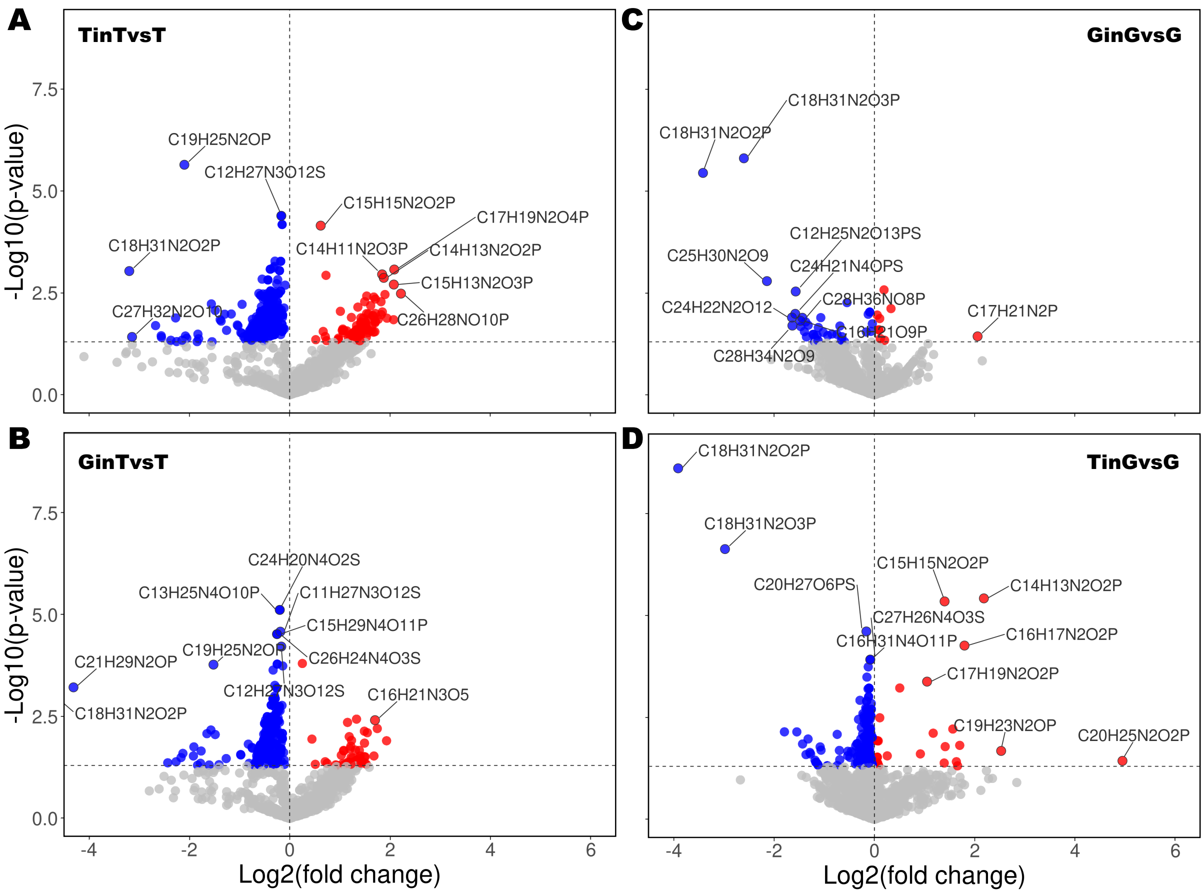


**Fig. S.I.1: Volcano plot of the molecules identifed by HRMS for each condition.** The volcano plot shows the fold-change (x-axis) versus the significance (y-axis) of the identified molecules in (**A**) TinT, (**B**) GinT, (**C**) GinG and (**D**) TinG conditions. The significance and the fold-change are converted to −Log10(p-value) and Log2(fold-change), respectively. The vertical and horizontal dotted lines show the cut-off of fold-change = 0, and of p-value = 0.05, respectively.

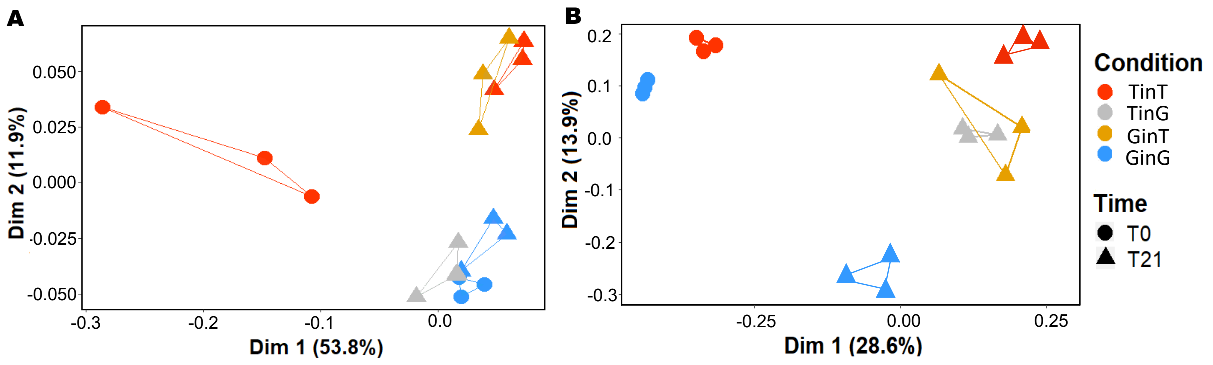


**Fig. S.I.2: Principal coordinate analysis (PCoA) plots derived from (A)** DOM molecular composition in each condition and at the beginning and the end of the incubation experiment and **(B)** from prokaryotic community composition in each condition and at the beginning, after 4 days of incubation and at the end of the incubation experiment.

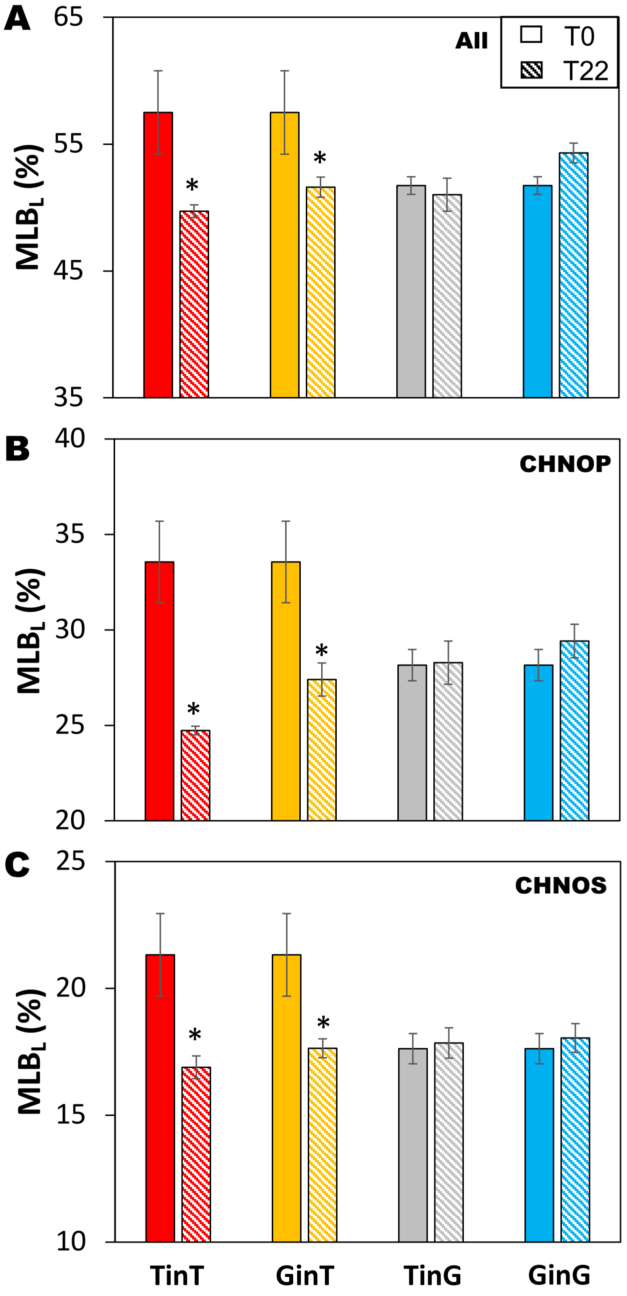


**Fig. S.I.3: Percentage of labile compounds of the dissolved organic matter in each condition at the initial (filled colors) and final (hatch) incubation time determined according to the molecular lability boundary (MLB)** among (A) all elementary formulas combined, (B) the heteroatomic molecular species containing carbon, hydrogen, nitrogen, oxygen and phosphate (CHNOP) and (C) the heteroatomic molecular species containing carbon, hydrogen, nitrogen, oxygen and sulphate (CHNOS). Error bars denote standard deviation of triplicate analyses.

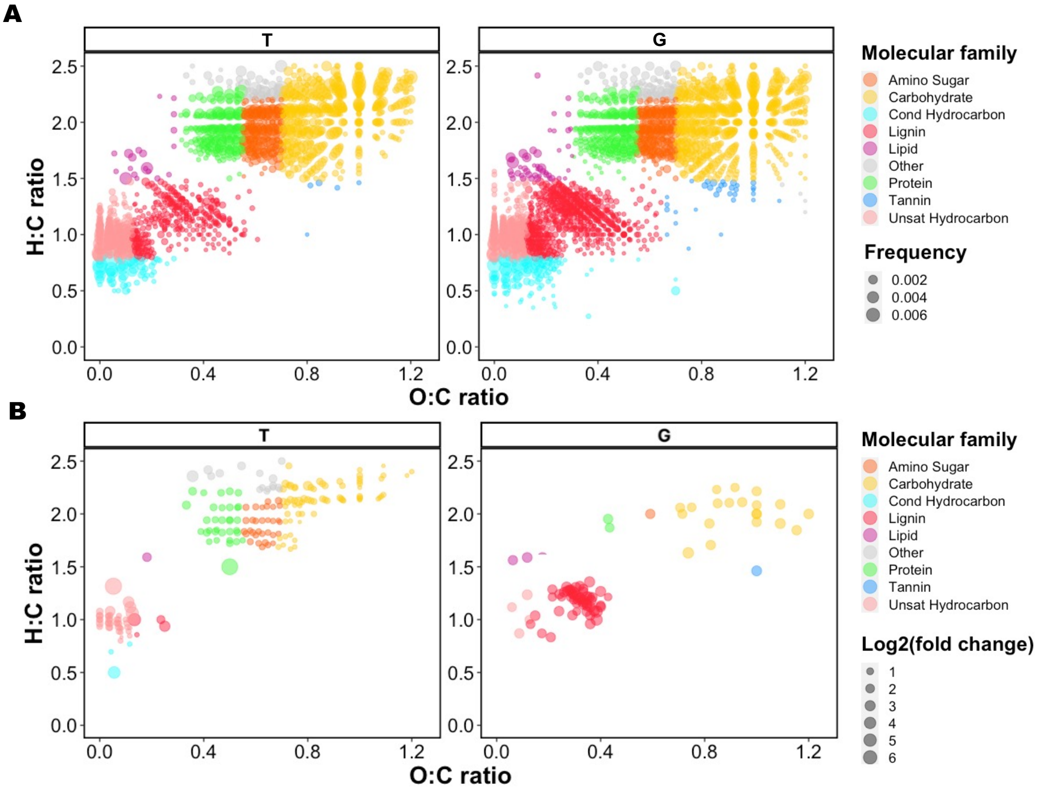


**Fig. S.I.4: Van Krevelen diagram showing (A) all the molecules composing dissolved organic matter from T and G sites. (B) The molecules composing dissolved organic matter allowing to explain the initial differences between the two sites.** The left panel represents the molecules preferentially present in T while the right panel represents the molecules preferentially present in G. The molecules are coloured according to their molecular family. The size of the dots is proportional to the variation factor of a molecule between the two sites.

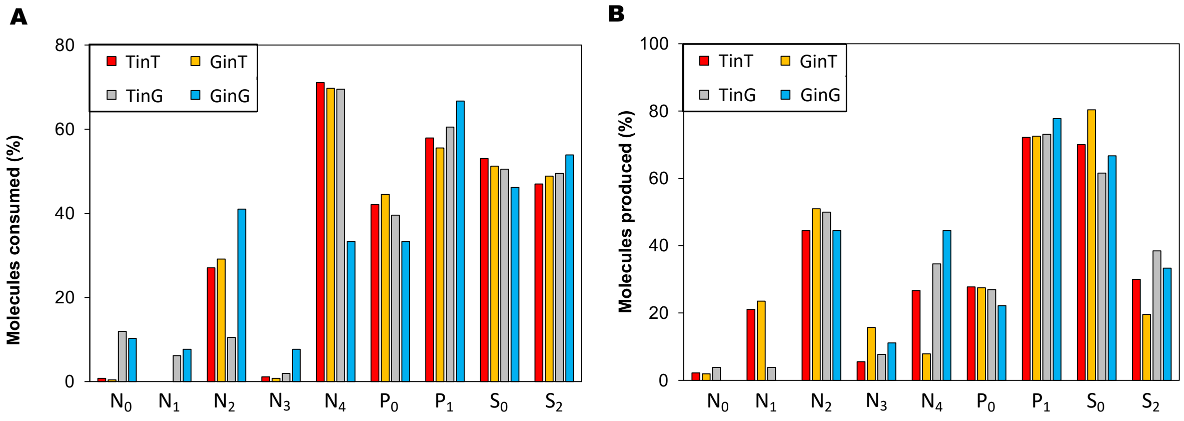


**Fig.S.I.5: Fraction of molecules (A) significantly consumed and (B) significantly produced according to the number of nitrate (N), phosphor (P) and sulfur (S) in their molecular formula for each experimental condition.**

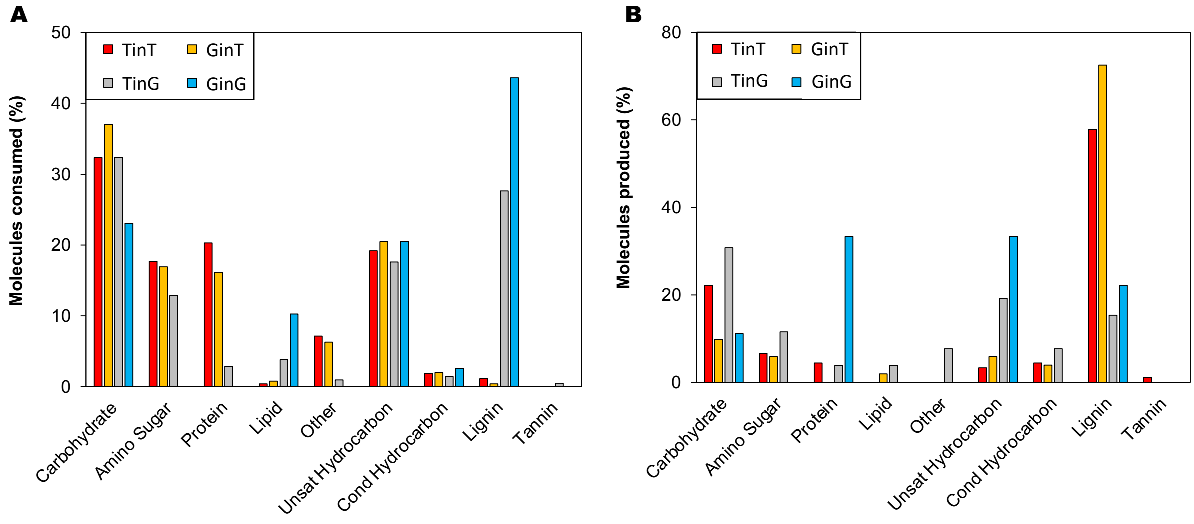


**Fig. S.I.6: Fraction of molecules (A) significantly consumed and (B) significantly produced according to the different molecular groups for each experimental condition.**
